# Epithelial-intrinsic nitric oxide synthase 2 sustains host-microbiota dynamics that promote colitis

**DOI:** 10.1101/2025.09.23.677925

**Authors:** Seika Hashimoto-Hill, Bryce Hunt, Sekani Cole, Shikha Negi, Amanda Waddell, Zi F. Yang, Joseph A. Wayman, Nicholas J. Ollberding, Emily R. Miraldi, Hee-Woong Lim, Andres Vazquez-Torres, Thomas E. Morrison, Lee A. Denson, Theresa Alenghat

**Author notes:** CORRESPONDENCE: Theresa Alenghat, 3333 Burnet Ave, MLC 7038.

## Abstract

The microbiota influence disease pathogenesis and treatment, however we have limited ability to assess patient status in relation to the microbiota. Here we find that the nitric oxide generating enzyme, nitric oxide synthase 2 (Nos2), is transcriptionally primed in intestinal epithelial cells (IECs), as opposed to immune cells, in inflammatory bowel disease (IBD) patients. Generation of IEC-specific Nos2 knockout mice revealed that epithelial Nos2 activity promoted susceptibility to intestinal disease and sustained a colitogenic microbiota. Epithelial Nos2 increased levels of nitric oxide-derived nitrates and nitrate-metabolizing bacteria in the intestine. Unexpectedly, extra-intestinal nitrates also reflected IEC-intrinsic Nos2 expression, and systemic nitrate concentrations in patients paralleled intestinal Nos2 activation. In fact, temporally inhibiting epithelial Nos2 was sufficient to alter intestinal nitrate homeostasis and inflammation in mice, as well as restrict nitrate production by human intestinal organoids. These data reveal that epithelial nitric oxide metabolism directs host-microbiota dynamics that can alter disease and that monitoring and targeting this axis may benefit patients with IBD.

## INTRODUCTION

Inflammatory bowel disease (IBD), including Crohn’s disease (CD) and ulcerative colitis (UC), affects over 5 million people globally and represents a significant public health and economic challenge^1^. Immune dysregulation that manifests in IBD patients as chronic inflammation is often characterized by stages of remission and relapse. Current therapeutics are directed towards decreasing excessive inflammation, rather than preventing early pathways that predispose to pro-inflammatory conditions. Although these treatments can ameliorate symptoms in many patients, side effects can be significant and treatment failure is common^2,3^. Therefore, improved understanding of the mechanisms that promote IBD development will enable development of personalized approaches to treat the disease and predict response to therapy. While genetic susceptibility is evident in IBD, it is also clear that external triggers are essential factors that impact the development and severity of IBD^4,5^. In addition to environmental contributors, such as diet and smoking, alterations in the composition of the intestinal microbiota have also been closely associated with clinical disease^6–8^. However, although the microbiota in IBD patients have been linked to disease pathogenesis, the mechanisms that sustain pathogenic microbiota that promote inflammation are not known.

Nitric oxide is a free radical synthesized from the amino acid L-arginine by nitric oxide synthases. Two nitric oxide synthase isoforms are constitutively present in either neuronal or endothelial tissue. A third isoform, termed inducible nitric oxide synthase or nitric oxide synthase 2 (Nos2), can be expressed at varying levels by immune cells, epithelial cells and endothelial cells^9^. Nos2 has been largely examined in macrophages, in which Nos2 expression is upregulated in response to pathogenic bacteria and inflammatory stimuli such as cytokines^9,10^. Nitric oxide has been described to exert multiple functions, including activation of guanylate cyclase, S-nitrosylation of protein thiols, and oxidative nitration^11–13^. Interestingly, Nos2 expression has been found to be highly increased in intestinal samples of both CD and UC patients across pediatric and adult patient cohorts^14,15^. However, *in vivo* studies of Nos2 in the context of intestinal inflammation have been conflicting^14–16^, likely because murine studies have relied on mice that lack Nos2 expression in all cells.

Intestinal epithelial cells (IECs) reside at the direct interface between the host and intestinal microbiota and IEC dysfunction characterizes IBD. IECs sense commensal bacterial-derived signals and coordinate host-microbiota symbiosis by maintaining barrier function and immune cell homeostasis in the intestine^17^. Here, we find that Nos2 transcriptional and epigenetic alterations in the intestine of IBD patients are unexpectedly restricted to IECs. Direct functional examination of epithelial-intrinsic Nos2 revealed that epithelial expression of this enzyme increased susceptibility to colitis. Epithelial-intrinsic Nos2 regulated intestinal concentrations of nitric oxide-derived nitrates and sustained elevated nitrate-metabolizing bacteria in the intestine that promote colitis. Similar to observations in adult patients^18,19^, nitric oxide derivatives in the blood and urine were elevated in pediatric IBD patients. Cell-specific murine models unexpectedly demonstrated that systemic nitrate levels highly reflected IEC-intrinsic Nos2 activity. In fact, short-term inhibition of epithelial Nos2 activity *in vivo* was sufficient to shift nitrate and microbial homeostasis, and improve colitis. Furthermore, nitrate regulation by Nos2 could effectively be targeted in patient-derived intestinal organoids. Collectively, these data indicate that priming of epithelial-intrinsic Nos2 may be a key mechanism that enables expansion of a colitogenic microbiota in IBD patients, and suggests that epithelial Nos2 activity may be clinically trackable and therapeutically targeted.

## RESULTS

### Nos2 expression is specifically elevated by intestinal epithelial cells of CD patients

Elevated Nos2 expression has been observed in intestinal samples collected from IBD patients^20–22^. To determine the cellular sources of Nos2 in the intestine of Crohn’s Disease (CD) patients, we analyzed single-cell RNA-sequencing datasets from ileal biopsies of treatment-naïve pediatric CD and non-IBD control patients^23^. Expression of Nos2 in non-IBD samples largely occurred in the epithelial population, whereas levels in immune cells, endothelial cells, and fibroblasts were below detection (**Figure 1A**). Furthermore, elevated Nos2 in biopsies from CD represented increased expression of this enzyme specifically in epithelial cells (**Figure 1A**). We therefore examined levels of the activating histone modification H3-lysine 4 trimethylation (H3K4me3) in the Nos2 gene in IECs purified from intestinal biopsies of treatment-naïve CD and non-IBD pediatric patients at Cincinnati Children’s Hospital^24^. Interestingly, H3K4me3 enrichment in the Nos2 promoter of IECs significantly correlated with expression of the IBD biomarker S100A8 (**Figure 1B**), suggesting a close link between epithelial Nos2 regulation and disease.

**Figure 1.**
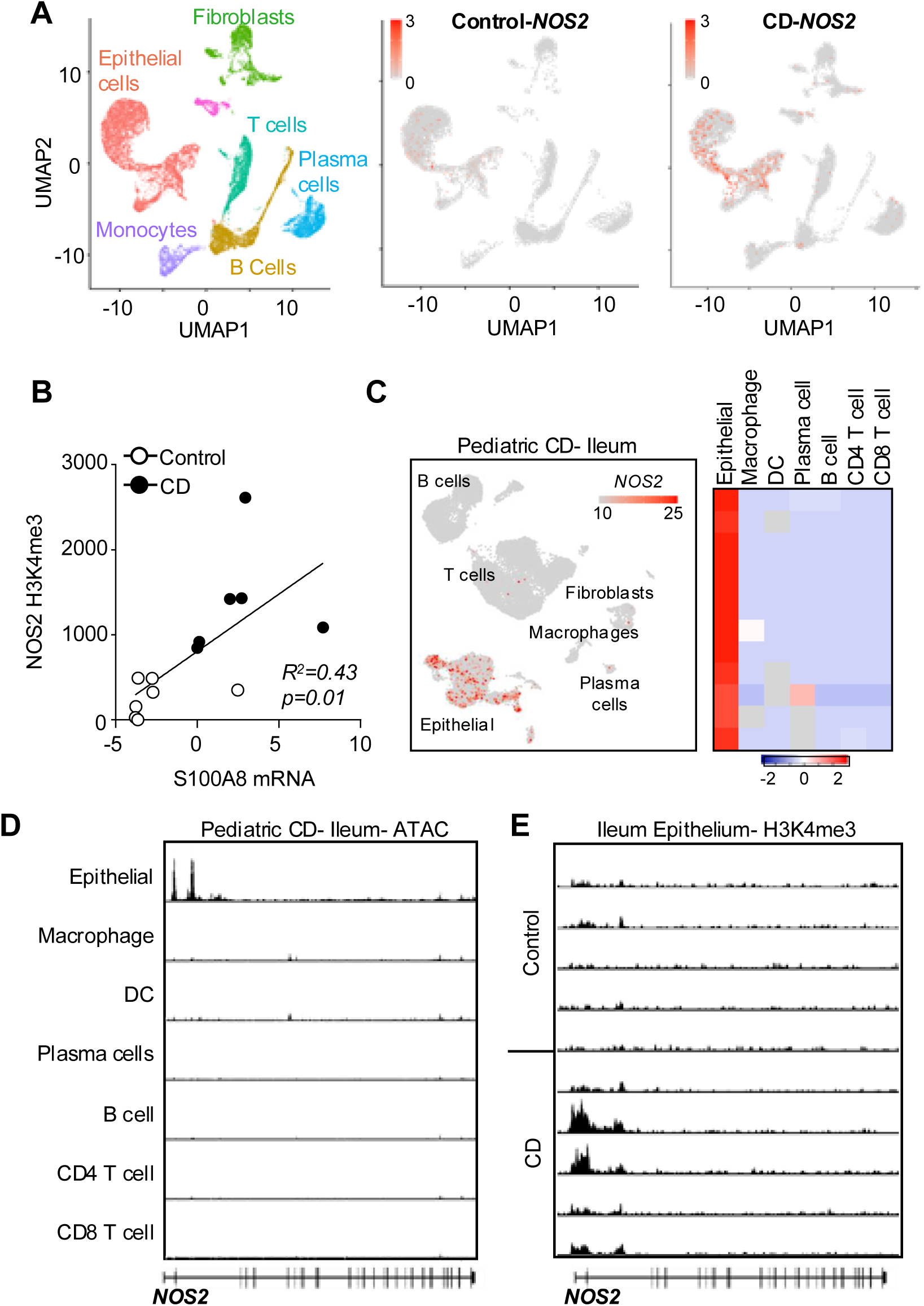
**Nos2 is specifically upregulated by epithelial cells in CD patients**. **(A)** Nos2 expression in single cell RNA-seq datasets from ileal biopsies of pediatric non-IBD (control) and Crohn’s disease (CD) patients (*21*). **(B)** Correlation between histone H3 Lysin4 tri-methylation (H3K4me3) enrichment by ChIP-seq in epithelial cells and expression of S100A8 from ileal biopsies of pediatric control and newly-diagnosed CD patients (*22*). **(C, D)** Sn-multiome-seq of ileal endoscopic biopsies from established pediatric CD patients. (C) Nos2 expression at single-cell resolution (left panel, log(transcripts per)) and donor-resolved pseudobulk resolution (right panel, z-scored DESeq2 VST counts, n=12). (D) Pseudobulk chromatin accessibility at the *NOS2* locus, normalized as fragments per million per cell type signal track. **(E)** ChIP-seq for H3K4me3 in epithelial cells isolated from ileal biopsies of a distinct pediatric cohort with control and established CD patients.

To examine cell-intrinsic expression and chromatin in parallel, we next performed single-nuclei (sn)multiome-seq (simultaneous snRNA-and snATAC-seq) analyses of ileal biopsies from a distinct pediatric cohort of CD patients. The same cellular expression profile was identified in which epithelial cells were the major producers of Nos2, relative other cell lineages in the intestine (**Figure 1C**). Chromatin analyses of these samples also demonstrated uniquely elevated accessibility at the Nos2 locus in epithelial cells (**Figure 1D**). We next harvested IECs from ileal endoscopic biopsies of CD patients in clinical remission and directly assessed H3K4me3 enrichment. H3K4me3 remained enriched, but varied in magnitude, in the Nos2 locus in CD samples relative to non-IBD controls processed together (**Figure 1E**), highlighting that transcriptional priming of IEC-intrinsic Nos2 represents a shared feature in the ileum of both treatment-naïve and established CD patients. Taken together, these patient findings reveal a robust transcriptional and epigenetic Nos2 signature in epithelial cells of IBD patients, and provoke the hypothesis that IEC-intrinsic regulation of Nos2 may have a functional role in calibrating susceptibility to intestinal inflammation.

### Epithelial *Nos2* expression increases susceptibility to microbiota-driven colitis

The epithelium-enriched expression of Nos2 in patients with IBD suggested a potential pathogenic role in inflammation. Therefore, to examine whether Nos2 contributes to intestinal inflammation in an epithelial-specific manner, we generated mice with IEC-specific deletion of Nos2 (Nos2^ΔIEC^) by crossing floxed-Nos2 mice (Nos2^FF^) with mice expressing Cre recombinase under the control of the Villin promoter (**Figure 2A**). IECs harvested from the intestine of Nos2^FF^ and Nos2^ΔIEC^ confirmed effective loss of Nos2 expression (**Figure 2B**). Nos2^FF^ and Nos2^ΔIEC^ mice were separated at weaning and subsequently subjected to the dextran sulfate sodium (DSS)-induced colitis model in adulthood. As expected, floxed control mice demonstrated significant weight loss (**Figure 2C**), colon shortening (**Figure 2D**), and severe disease (**Figure 2E**), following DSS administration. However, mice lacking IEC-specific expression of Nos2 exhibited significantly reduced susceptibility to colitis, as evidenced by attenuated weight loss (**Figure 2C**), colon shortening (**Figure 2D**) and disease score (**Figure 2E**). Consistent with these clinical parameters, histologic examination of the colon was characterized by ulceration, edema, and inflammatory infiltration in Nos2-expresing controls, whereas these pathologic features of colitis were markedly diminished in DSS-exposed Nos2^ΔIEC^ mice (**Figure 2F**).

**Figure 2.**
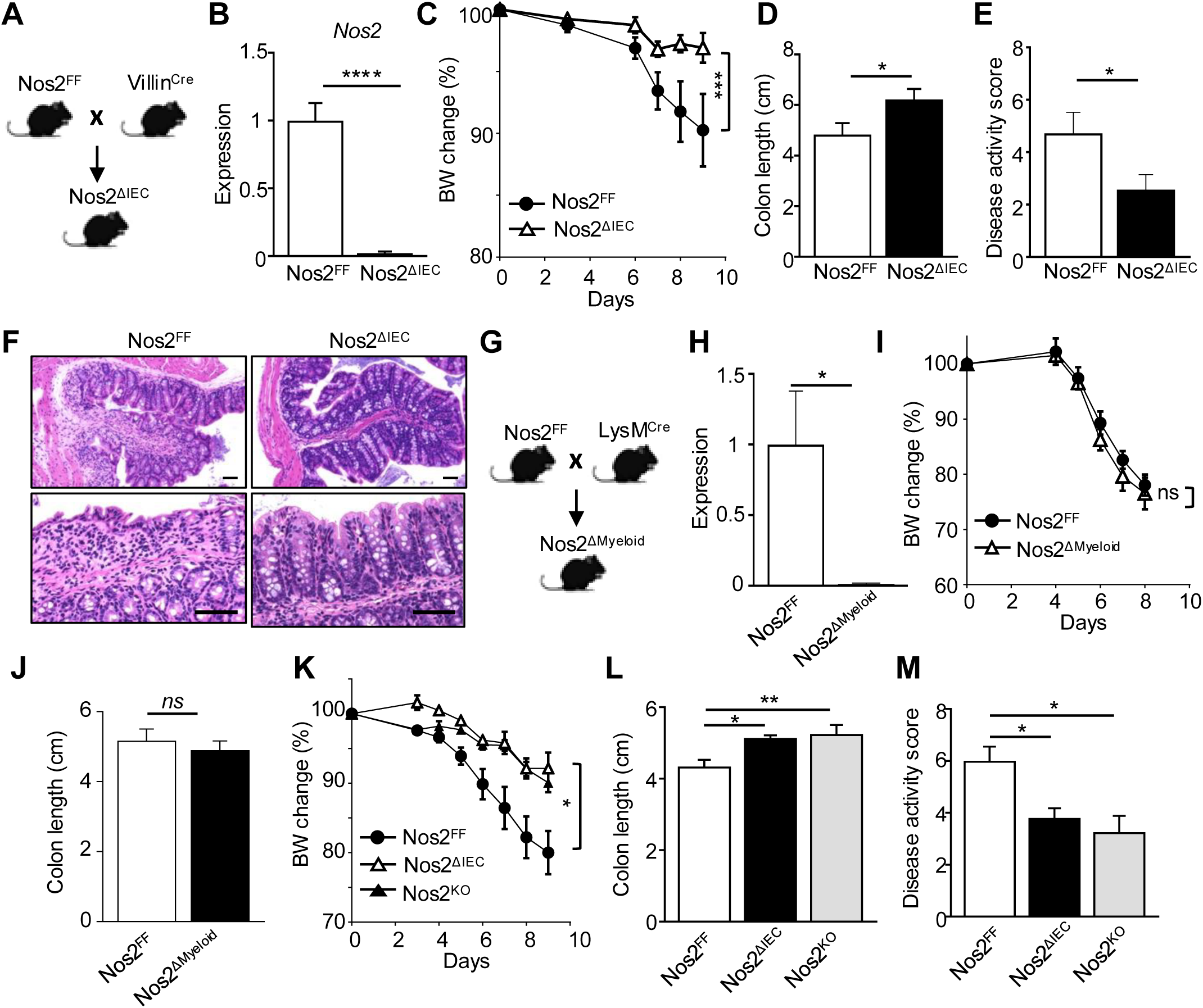
Epithelial-intrinsic *Nos2* expression increases susceptibility to colitis. (A) Approach to generate IEC-specific deletion of Nos2 in mice (Nos2^ΔIEC^) by breeding floxed (Nos2^FF^) mice to mice expressing cre recombinase downstream of the villin promoter (Villin^Cre^). **(B)** Nos2^ΔIEC^ mRNA expression in LI IECs harvested from Nos2^FF^ mice relative to Nos2 mice. **(C)** Body weight (BW) changes due to DSS-induced colitis. Nos2^FF^ mice and Nos2^ΔIEC^ mice were generated in the same breedings then separately housed based on gender and genotype at weaning. Mice received 3% DSS in their drinking water for 6-7 days, and the returned to regular water for the remainer of study. **(D)** Colon length, **(E)** Disease activity and **(F)** H&E-stained colon on day 9 following DSS, scale bars 50 μm. **(G)** Mouse model of myeloid-specific deletion of Nos2. **(H)** Nos2 mRNA expression in peritoneal macrophages. **(I)** BW changes due to DSS colitis. **(J)** Colon length on day 8 following DSS. **(K)** BW changes of Nos2FF, Nos2^ΔIEC^ and total Nos2 knockout (Nos2^KO^) due to DSS colitis. **(L)** Colon length **(M)** Disease activity on day 9 following DSS. Data are representative of at least three independent experiments, 3-4 mice per group. Results are mean ± SEM. * p<0.05, ** p<0.01, ***p<0.001.

The regulation and function of Nos2 have been extensively characterized in murine macrophages, where its pro-inflammatory role has been implicated in intestinal inflammation^9,10,25,26^. However, the necessity of Nos2 expression in myeloid cells for the development of intestinal inflammation has not been systematically investigated *in vivo* using cell-specific conditional knockout mice. To address this, we generated mice with myeloid-specific deletion of Nos2 (Nos2^ΔMyeloid^) by crossing floxed-*Nos2* (Nos2^FF^) mice with a LysM-Cre transgenic line (**Figure 2G**). Deletion of Nos2 in macrophages was confirmed in Nos2^ΔMyeloid^ mice (**Figure 2H**), and Nos2^FF^ and Nos2^ΔMyeloid^ mice were then subjected to DSS-induced colitis to assess the role of macrophage-expressed Nos2 in disease pathogenesis. In contrast to the significant reduction in disease susceptibility observed in Nos2^ΔIEC^ mice (**Figure 2C-F**), Nos2^ΔMyeloid^ mice exhibited comparable susceptibility to floxed Nos2 controls (**Figure 2I, J**). These findings indicate that epithelial-expressed Nos2 exerts a pathogenic role in DSS-induced intestinal inflammation, whereas myeloid-derived Nos2 does not contribute significantly to disease progression. Previous studies reported that global Nos2 knockout (Nos2^KO^) mice are protected from DSS-induced colitis, attributing loss of Nos2 in myeloid cells as the principal cell responsible^27–29^. However, direct comparison of Nos2^FF^, Nos2^ΔIEC^, and Nos2^KO^ mice demonstrated that Nos2^ΔIEC^ phenocopy parameters observed in Nos2^KO^ mice when subjected to DSS (**Figure 2K-M**). These data support that the phenotype observed in Nos2^KO^ mice is largely attributable to the loss of epithelial-intrinsic *Nos2* expression.

### Loss of *Nos2* in IECs decreases colonization with colitogenic bacteria

Colitis is influenced by variations in the gut microbiota^30–32^. Moreover, host-derived nitric oxide can modulate microbiota composition during inflammation by selectively promoting the growth of bacteria capable of nitrate respiration^33,34^. To assess whether microbiota contribute to the Nos2-dependent DSS phenotype, littermate Nos2^FF^ and Nos2^ΔIEC^ mice remained cohoused to normalize for differences in microbiota. The mice were then subjected to DSS-induced colitis. Interestingly, Nos2^ΔIEC^ mice were not protected from DSS-induced colitis when cohoused with Nos2-replete littermate control mice (**Figure 3A**), suggesting that the commensal microbes may be sensitive to epithelial-intrinsic Nos2 activity in the intestine. To investigate this, we employed metagenomic analyses to broadly compare the composition of the intestinal microbiota at steady state in 8 week-old Nos2^FF^ and Nos2^ΔIEC^ littermate mice that were separated at weaning. Significant differences were not identified at the phyla level (**Figure S1A**), and the alpha-diversity as measured by the Shannon Index was similar between Nos2^FF^ and Nos2^ΔIEC^ mice (**Figure S1B**). Nitric oxide produced by Nos2 is converted to nitrates that can be utilized for respiration by specific commensal bacterial strains^16^. Notably, we found that these nitric oxide derivatives were decreased in the colonic mucosa of Nos2^ΔIEC^ mice compared to Nos2^FF^ controls (**Figure 3B**). Based on this observation, we directly examined the abundance of nitrate-utilizing bacterial species. Interestingly, loss of Nos2 expression in IECs led to a reduction in multiple known nitrate-metabolizing bacterial families, including Desulfovibrionaceae, Helicobacteraceae, Mucispirillaceae, Lactobacillaceae, and Burkholderiaceae (**Figure 3C**).

**Figure 3.**
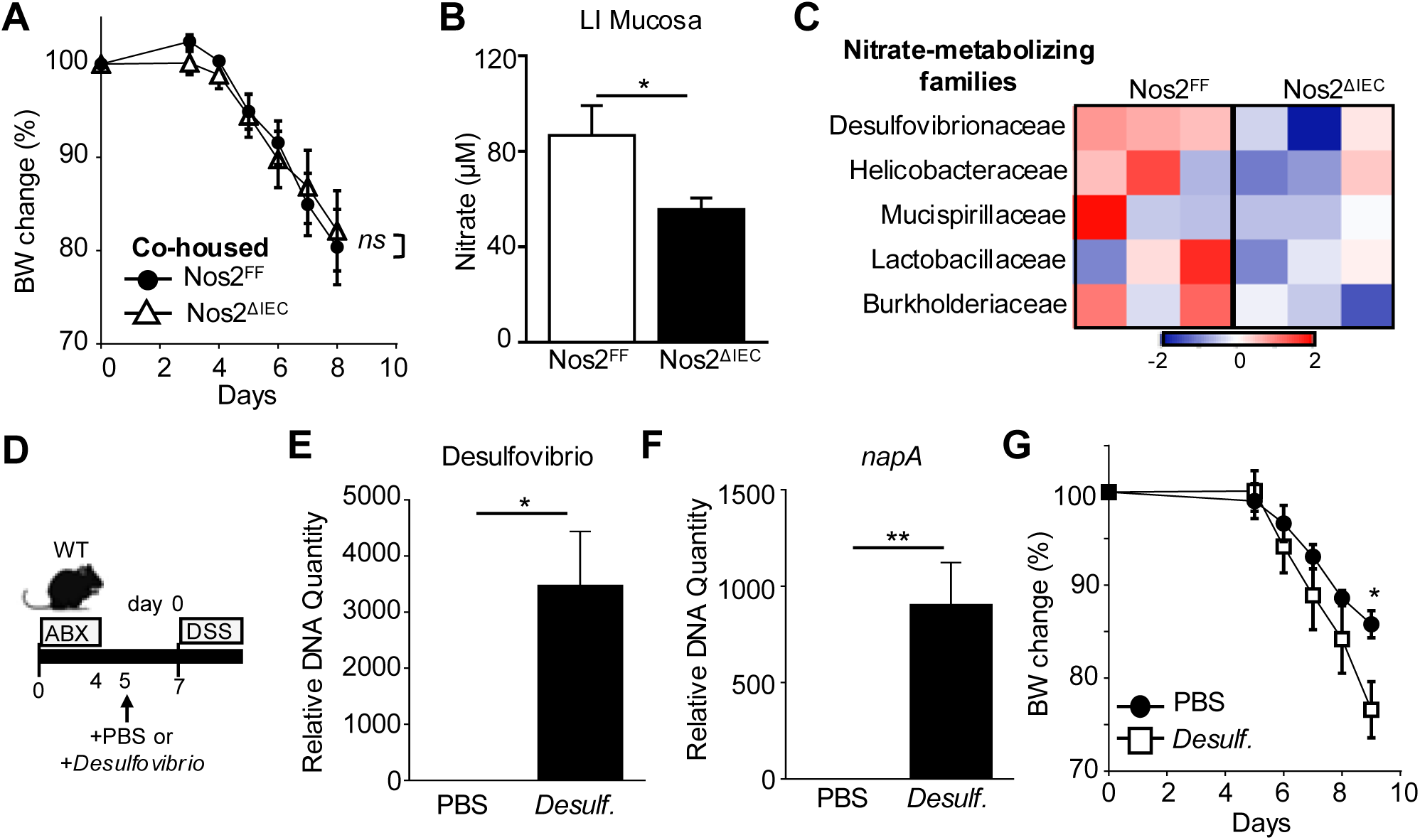
Loss of *Nos2* in IECs decreases colonization with colitogenic bacteria. (A) **BW** changes due to DSS colitis in cohoused Nos2^FF^ mice and Nos2^ΔIEC^ mice. **(B)** Nitrate concentrations measured by modified Griess assay in mucosal scrapes from the large intestine of naïve Nos2^FF^ mice and Nos2^ΔIEC^ mice separated by genotype at weaning. **(C)** Relative abundance of nitrate-utilizing bacterial families based on metagenomic analyses of stool samples, shown as z-scores. **(D)** Experimental design to colonize wildtype mice with Desulfovibrio desulfuricans. Antibiotics (ABX) were administered for 4 days prior to gavage with Desulfovibrio desulfuricans. **(E, F)** (E) Desulfovibrio 16S and (F) the nitrate reductase gene, napA, abundance in stool prior to DSS, measured by qPCR. **(G)** BW changes due to DSS colitis. Data are representative of at least three independent experiments, 3-4 mice per group. Results are mean ± SEM. * p<0.05, ** p<0.01.

An increased abundance of nitrate-respiring bacteria has been associated with intestinal inflammation and IBD, and suggested to promote more severe intestinal inflammation^33–35^. Desulfovibrionaceae is a nitrate-utilizing commensal that demonstrated enrichment in mice expressing epithelial Nos2, relative to Nos2-deficient mice. Thus, to assess how the presence of these bacteria in Nos2-replete mice impacts development of colitis, we employed antibiotics to create an open niche and then colonized with Desulfovibrio desulfuricans^36^ (**Figure 3D**). Effective Desulfovibrio colonization was confirmed by measuring the Desulfovibrio 16S rRNA gene (**Figure 3E**) and napA, the gene encoding periplasmic nitrate reductase in Desulfovibrio (**Figure 3F**). Following exposure to DSS, Desulfovibrio-colonized mice developed more severe colitis relative to mice lacking Desulfovibrio (**Figure 3G**). Collectively, these findings indicate that epithelial Nos2 activity in the intestine increases mucosal concentration of nitrates and enable colonization with colitogenic nitrate-respiring bacterial populations.

### Nitrates represent a systemic biomarker of intestinal epithelial NOS2 activity

To examine whether Nos2 could be targeted to improve susceptibility to intestinal damage and inflammation, we employed the Nos2 inhibitor aminoguanidine (AMG). Similar to Nos2^ΔIEC^ and Nos2^KO^ mice, administration of AMG attenuated colitis in mice following DSS exposure (**Figure 4A-C**). Nitric oxide is rapidly oxidized to nitrate and AMG treatment led to a measurable decrease in systemic nitrates (**Figure 4D**). Remarkably, mice lacking IEC-intrinsic Nos2 expression also exhibited a marked decrease in serum (**Figure 4E**) and urine (**Figure 4F**) nitrate levels compared to Nos2-replete floxed mice (**Figure 4E, F**). The levels in Nos2^ΔIEC^ mice were in fact more comparable to full Nos2^KO^ mice that lack Nos2 expression in all cell lineages (**Figure 4E, F**). In addition, nitrate levels in mice lacking Nos2 only in myeloid cells were similar to controls (**Figure 4G, H**). Taken together, these data support that systemic nitrate levels are highly reflective of IEC-intrinsic Nos2 expression. Interestingly, administration of AMG to mice reduced urinary nitrate concentrations to levels comparable to those observed in Nos2^ΔIEC^ mice (**Figure 4I**). Furthermore, AMG-treated Nos2^ΔIEC^ mice did not exhibit a further reduction in nitrate levels relative to Nos2^ΔIEC^ mice that did not receive AMG (**Figure 4I**), indicating that AMG-mediated inhibition of nitrate levels requires epithelial Nos2 activity.

**Figure 4.**
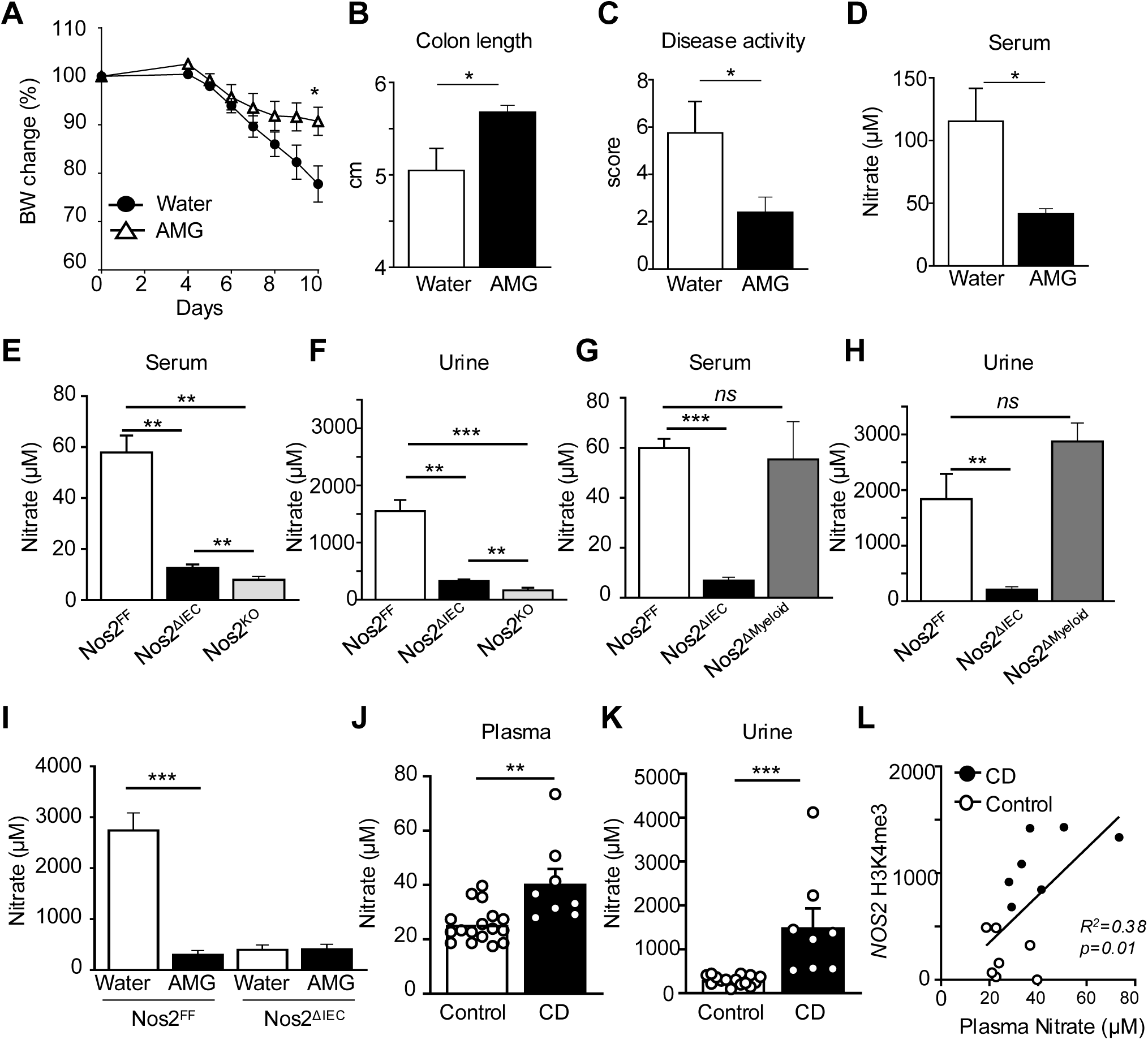
Nitrates represent a systemic biomarker of intestinal epithelial NOS2 activity. **(A)** BW changes due to DSS colitis in wildtype mice receiving the Nos2 inhibitor aminoguanidine (AMG) for 1 week prior to and during DSS administration. **(B)** Colon length and **(C)** Disease activity score for mice in (A) on day 10 following DSS. **(D)** Nitrate concentrations measured by modified Griess assay in the serum of DSS treated wildtype mice following 1 week of receiving 1% AMG in their water. **(E-H)** Nitrate concentrations in (E, G) serum and (F, H) urine of naive mice. **(I)** Nitrate concentrations in the urine of vehicle and AMG-treated Nos2^FF^ and Nos2^ΔIEC^ mice. Data are representative of at least three independent experiments, 3-4 mice per group. **(J, K)** Nitrate concentrations measured by modified Griess assay in (J) plasma and (K) urine of non-IBD (control) and CD patients. **(L)** Correlation between patient plasma nitrate concentration and H3K4me3 enrichment by ChIP-seq in epithelial cells from patient ileal biopsies. Results are mean ± SEM. ** p<0.01, *** p<0.001, ns: not significant.

Elevated nitrate levels in blood and urine have been observed in adult patients with IBD^18,19,37^. Interestingly, we found that nitrate concentrations in both the plasma (**Figure 4J**) and urine (**Figure 4K**) of pediatric CD and non-IBD patients were also significantly higher in patients with CD compared to non-inflamed control patients (**Figure 4J, K**). Furthermore, serum nitrate concentrations positively correlated with H3K4me3 enrichment at the Nos2 gene in patient IECs (**Figure 4L**). While the underlying biochemistry for these extra-intestinal dynamics is not clear, this finding suggests that systemic nitrate levels in fact parallels Nos2 regulation in the intestine. Thus, clinically monitoring and pharmacologically targeting epithelial-intrinsic Nos2 activity may be beneficial for managing IBD.

### Nos2 activity in IECs actively directs microbiota and primes susceptibility to colitis

To test the temporal sensitivity of nitrate dynamics on IEC-intrinsic Nos2 and to eliminate potential developmental effects of constitutive deletion, we generated an inducible tamoxifen-dependent IEC-specific Nos2 knockout mouse model (Nos2^ΔIEC-IND^) (**Figure 5A**) in which depletion of Nos2 in IECs could be detected post tamoxifen treatment (**Figure 5B**). Remarkably, urine nitrate concentrations were already markedly lower in Nos2^ΔIEC-IND^ mice within 3 days following tamoxifen administration (**Figure 5C**), confirming the high sensitivity of systemic nitrate levels to IEC-intrinsic Nos2 regulation. Furthermore, the bacterial nitrate reductase gene napA decreased in the intestine of Nos2^ΔIEC-IND^ mice post tamoxifen exposure. Thus, epithelial Nos2 activity dynamically regulates the nitrate-metabolizing capacity of the intestinal microbiota (**Figure 5D**). Lasty, tamoxifen-treated Nos2^ΔIEC-IND^ mice subjected to DSS (**Figure 5E**) exhibited milder weight loss (**Figure 5F**), decreased colonic inflammatory infiltration (**Figure 5F-H**), and less severe intestinal pathology relative to control mice (**Figure 5I**). Collectively, these results identify a central role for epithelial-intrinsic Nos2 in actively regulating host-microbiota dynamics, susceptibility to colitis, and levels of measurable nitrates.

**Figure 5.**
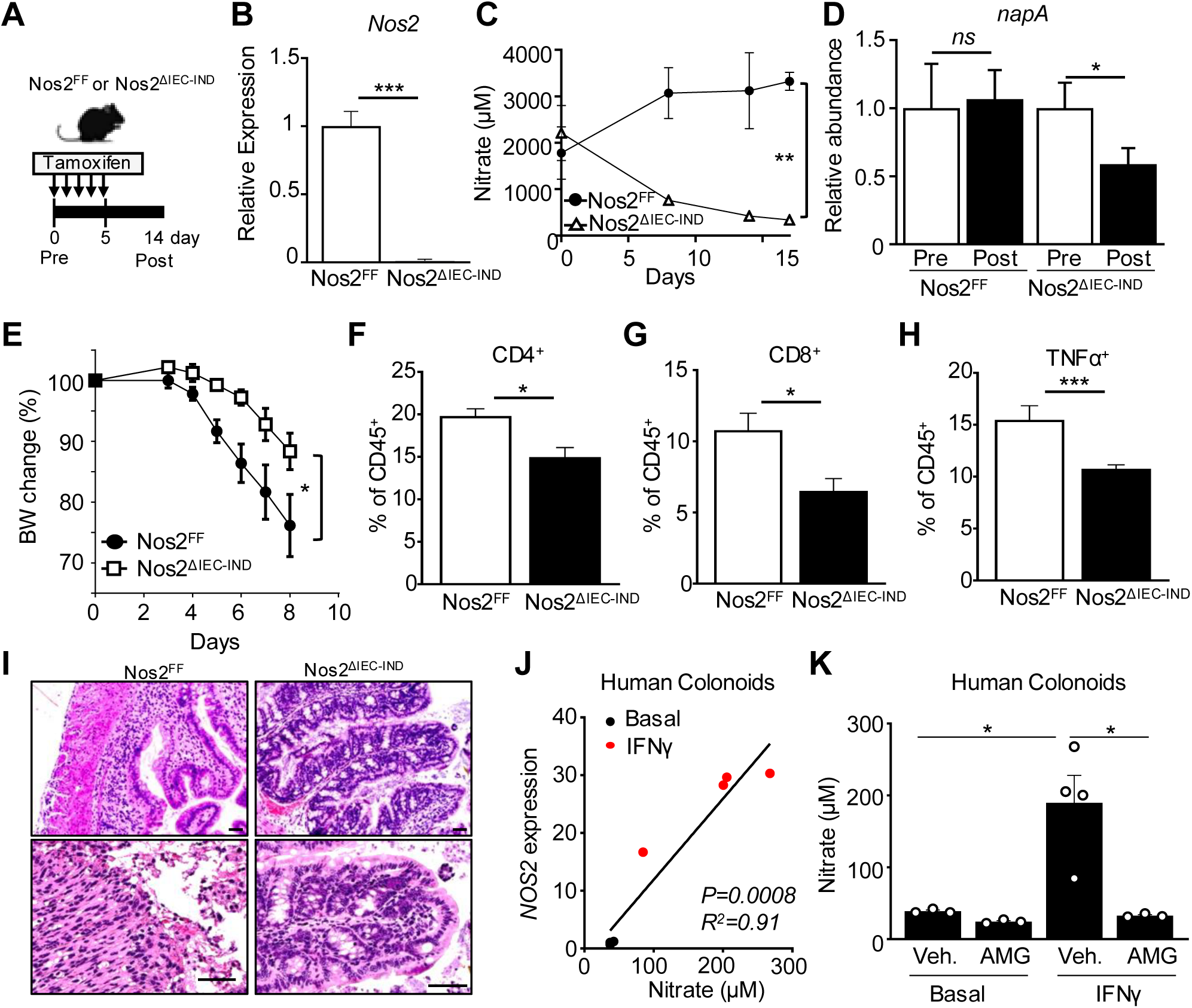
Nos2 activity in IECs actively directs intestinal nitrate metabolism and primes susceptibility to colitis. **(A)** Experimental design using an inducible tamoxifen-sensitive Nos2^ΔIEC-IND^ mouse model. Adult Nos2^FF^ and Nos2^ΔIEC-IND^ mice were treated with tamoxifen once a day for 5 consecutive days to induce loss of Nos2 in IECs. **(B)** Nos2 mRNA expression in LI IECs harvested from Nos2^ΔIEC-IND^ mice, relative to Nos2^FF^ mice 14 days after final dose of tamoxifen. **(C)** Temporal tracking of nitrate levels in urine following tamoxifen administration to Nos2^FF^ and Nos2^ΔIEC-IND^ mice. **(D)** Levels of the nitrate reductase gene napA by stool qPCR prior to and post tamoxifen. **(E)** BW changes of induced Nos2^FF^ or Nos2^ΔIEC-IND^ subjected to DSS. **(F-H)** Frequency of (F) CD4^+^ T cells, (G) CD8^+^ T cells, and (H) TNFα^+^ leukocytes in the colonic lamina propria of DSS-treated Nos2^FF^ and Nos2^ΔIEC-IND^ mice, gated on live, CD45. **(I)** H&E-stained colon tissues on day 8 following DSS, scale bars 50 μm. 3-4 mice per group. **(J)** Correlation between Nos2 expression and nitrate concentration in supernatant of human colon crypt-derived intestinal organoids (colonoids)-/+ IFNγ (50ng/ml). **(K)** Nitrate concentration in supernatant of human colonoids-/+ AMG (5mM). Colonoids were generated from two different patients. Results are mean ± SEM. * p<0.05, ** p<0.01, *ns*: not significant.

To translate the potential functional and therapeutic relevance of these findings in human intestine, we generated intestinal epithelial organoids (colonoids) from biopsies collected from the colon of control non-IBD patients. Human colonoids upregulated Nos2 expression in response to the cytokine interferon-gamma (IFNγ) (**Figure 5J**). Interestingly, Nos2 expression positively correlated with nitrate levels measured in the colonoid supernatant, suggesting that extracellular nitrate concentrations reflect human intestinal epithelial Nos2 activity (**Figure 5J**). Furthermore, similar to findings in mice, treatment of human colonoids with AMG blocked nitrate induction in an epithelial Nos2-dependent manner (**Figure 5K**). These human organoid data support that Nos2 activity in human intestinal epithelium actively modulates local nitrate concentrations and that this pathway could be evaluated therapeutically in patients.

## DISCUSSION

In this study, Nos2 was identified as an intrinsic factor of epithelial cells in the intestine of CD patients that promotes increased levels of nitrate-metabolizing bacteria and increases susceptibility to intestinal inflammation (**Figure S2**). Contrary to the prevailing focus on Nos2 expression in intestinal immune cells, our single cell analyses of intestine from CD patients revealed heightened Nos2 expression specifically in IECs. Interestingly, recent scRNA-seq analyses suggest that this could reflect an epithelial subtype present in the ileum of adult CD patient^38^. Further, epithelial Nos2 expression significantly influenced nitrate concentrations in blood and urine. Thus systemic nitrates can reflect the intestinal nitrate metabolic status of patients. Nitric oxide, a product of Nos2-mediated arginine metabolism, acts as a signaling molecule in various physiological processes in a dose-dependent manner. Nos2 generates elevated nitric oxide levels, which modulate cellular functions via guanylate cyclase activation and post-translational modifications of proteins, including S-nitrosylation, oxidative nitration, hydroxylation, and metal nitrosylation of transition metals^11–13^. Thus, nitric oxide has been speculated to exacerbate inflammation in IBD via direct action on mammalian cells^14,15^. Our findings add another layer of complexity to the disease pathogenesis, demonstrating that epithelial-derived nitric oxide promotes colonization with commensal bacterial strains that promote colitis (**Figure S2**).

Studies with epithelial cancer cell lines have suggested apical location of Nos2 and nitric oxide^39,40^. Interestingly, our human intestinal organoid analyses demonstrated that extracellular nitrate concentrations can be directly modulated by epithelial Nos2 in human intestine. Nitric oxide, although short-lived, exerts antimicrobial effects by producing reactive nitrogen species that target microbial components such as heme, iron-sulfur proteins, and DNA, inducing nitrosative stress^41,42^. Conversely, stable nitric oxide products like nitrate are utilized by certain bacteria for growth through nitrate respiration. Mice lacking IEC-intrinsic Nos2 expression were characterized by an intestinal microbiota with a reduced nitrate-metabolizing capacity. This aligns with findings from large scale-omics studies of IBD patients, which reported enrichment of nitrate reductase genes in dysbiotic CD metagenomes, as well as elevated Nos2 expression in the ileum^6^. Prior work found that host-derived nitric oxide promoted growth of commensal Enterobacteriaceae during intestinal inflammation^34^. This study conceptually advances our understanding of host-microbiota communication by demonstrating that epithelial-derived nitric oxide broadly influences microbiota composition even before inflammation onset, thereby setting up a host environment that predisposes to intestinal inflammation (**Figure S2**).

While elevated systemic nitrate levels have been observed in adult IBD patients^18,19,37^, the cellular source was not known, but presumed to largely reflect expression by inflammatory cell populations^37^. Unexpectedly, our data using constitutive and inducible mouse models demonstrated that systemic nitrate concentrations are instead temporally reflective of IEC-intrinsic Nos2 expression. Although it is not clear whether this metabolite sensitivity is a direct and/or indirect effect of intestinal Nos2 activity, systemic nitrate levels highly paralleled IEC-intrinsic Nos2 expression. Thus, systemic nitrate measurements may serve as a non-invasive tool to track a patient’s dynamic host-microbiota relationship in the context of health as well as disease, complementing conventional inflammatory biomarkers like calprotectin. Importantly patient analyses show conservation of the epithelial Nos2 axis in patient biopsies and human intestinal organoids, suggesting similar nitrate dynamics across species.

These findings identify epithelial Nos2 as a pivotal factor perpetuating host-microbiota interactions that can promote intestinal inflammation in IBD. Furthermore, we found that both human and murine intestinal epithelial Nos2 activity can be pharmacologically inhibited, suggesting that targeting this axis holds therapeutic potential for managing dysbiosis and intestinal inflammation in IBD. Despite past considerations of Nos2 inhibition, clinical trials have not yet been undertaken, possibly due to safety concerns over long-term inhibition^43^. While cytokines can influence Nos2 expression^44,45^, future studies that identify microbial factors in patients that initiate a transcriptionally active Nos2 state in IECs, may unveil additional approaches for calibrating epithelial Nos2 in the intestine. Developing targeted drug delivery systems for Nos2 inhibitors to epithelial cells could also enhance therapeutic specificity, particularly when combined with microbiota-based therapies. Collectively, these data reveal that diagnostically monitoring and targeting epithelial-intrinsic regulation of nitric oxide therapeutically holds promise for improving personalized treatment strategies in the management of IBD.

## MATERIALS AND METHODS

### Patients

Children and adolescents undergoing colonoscopy were recruited prospectively for participation in the IBD biorepository protocol between 2015 and 2021 at CCHMC. Adolescents, between the ages of 12 and 18 years, with treatment naïve Crohn’s disease or previously diagnosed CD were selected for this study, along with age-and sex-matched controls. Control patients presented due to intestinal symptoms, and colonoscopy was warranted as part of their evaluation. These patients were found to have no evidence of intestinal disease based on a combination of endoscopy, histology, lab testing, and imaging. Molecular analyses were performed on pooled ileal biopsy samples. A pediatric urine collection bag (Fisher cat. #22275347 or Hollister U-bag #7501) was used to collect their urine. Samples were stored in-80°C. Studies were performed with approval by the CCHMC Institutional Review Board (IRB 2011-2285).

**Animals.** Mice were housed up to 4 per cage in a ventilated isolator cage system in a 12-hour-light/dark cycle, with free access to water and chow. All experiments were performed with age-and sex-matched C57BL/6 mice. WT mice were obtained from The Jackson Laboratory. Nos2-/-mice were bred on site. Mice containing flox sites spanning exon 14-17 of Nos2 (Nos2^FF^) were crossed to villin-Cre-recombinase expressing mice to generate intestinal epithelial cell specific Nos2 knockout mice. Inducible IEC-specific Nos2 KO were generated by crossing Nos2^FF^ to tamoxifen-inducible Villin-Cre mice, then administering 1mg of tamoxifen i.p. once a day for 5 consecutive days. LysM-Cre-recombinase was used to generate Nos2^ΔMyeloid^ mice. Desulfovibrio *desulfuricans* subsp. *Desulfuricans* ATCC 27774 was obtained from DSMZ and anaerobically cultured in ATCC medium 1249 modified Barr media. Mice were treated with streptomycin (5mg/ml), ampicillin (1mg/ml), colistin (1mg/ml), metronidazole (1mg/ml) and sucralose (1mg/ml) for 4 days prior to Desulfovibrio colonization. Antibiotics were discontinued 24hrs prior to Desulfovibrio gavage. The Nos2 inhibitor aminoguanidine was added to drinking water at the concentrations of 0.1 or 0.2%. All murine experiments were performed according to the animal experimental guidelines upon approval of the Institutional Animal Care and Use Committee at CCHMC.

### Single-cell RNA and single nuclei multiome analyses

Ileal biopsy single-cell RNA sequencing from pediatric Crohn’s Disease and matched healthy patient datasets were downloaded from CZ CELL x GENE portal (https://cellxgene.cziscience.com/e/8e47ed12-c658-4252-b126-381df8d52a3d.cxg)^23^. The data were reprocessed using Seurat from variable gene identification, principal component analysis, UMAP projections, and clustering. NOS2 (ENSG00000007171) expression was project onto the UMAP using Seurat. Single nuclei multiome of terminal ileal biopsies were processed as described^46^. These data are available from GEO Database accession number GSE274451.

### IEC isolation

For isolation of IECs, samples were incubated with 1 mM EDTA and 2 mM DTT while shaking at 37°C at a 45° angle for 10 minutes to obtain high epithelial purity with similarly small amounts of CD45^+^ cells (≤5%) in both control and CD samples. Supernatant with IECs was pipetted into a new 50-ml tube and pelleted at 500 *g* for 5 minutes at 4°C. A portion of IECs was processed for ChIP-seq. Mouse IECs were harvested from 10 cm of the distal small intestine or large intestine. The intestine was opened longitudinally, washed, and incubated in buffer containing 1 mM EDTA, 2 mM DTT, and 5% FBS in PBS for 10 minutes, while shaking at 37°C at a 45° angle to isolate IECs.

### ChIP-seq

Fresh formaldehyde was added to each IEC sample for a final concentration of 1% and mixed end over end at room temperature for 10 minutes to crosslink proteins to DNA. Cells were lysed with a Triton X-100 and Igepal buffer (0.25% Triton C-100, 0.5% Igepal, 10% glycerol, 1 mM EDTA, 140 mM NaCl, and 50 mM HEPES). Nuclei were then washed and chromatin sheared to 150-to 500-bp size in a 0.1% SDS in TE buffer using a S220 Covaris Sonicator.

Immunoprecipitation was carried out with a SX-8G IP-STAR robot (Diagenode) and an antibody optimized for H3K4me3. Sequencing was performed using an Illumina HiSeq 2500. S100A8 expression correlation was assessed by RNA-sequencing as described previously^24^. ChIP-seq sequencing reads were aligned to USCS human hg38 using a STAR aligner^47^. Uniquely aligned reads were retained for downstream analysis. Aligned reads were down sampled to 30 million. BigWig files were generated using bedtools^48^ and UCSC toolkit in a read-per-million (RPM) scale.

### Real-time PCR

RNA from primary IECs and organoids were isolated using the RNeasy Kit (Qiagen) following manufacturer’s protocol. RNA was reverse-transcribed with Verso reverse transcriptase (Invitrogen) and expression was compared using SYBR (Applied Biosystems) and analyzed in the linear range of amplification. For quantification of bacterial genes in stool DNA, 4.5ng of DNA was used per PCR reaction. Primer sets were Desulfovibrio 16S 5-CCGTAGATATCTGGAGGAACATCAG-3, 5-ACATCTAGCATCCATCGTTTACAGC-3^49^, and NapA 5-AAGGAAAGGGTCACACAGCC-3, 5-CATCCCAGCTCACAGGTTCA-3.

### Shotgun metagenome sequencing and analyses

Fecal pellets were collected and stored in - 80C. Samples were transferred to barcode collection tubes containing DNA stabilization buffer (Transnetyx, Cordova, USA). DNA extraction, library preparation, and shotgun metagenomic sequencing were performed by TransnetYX (Cordova, TN, USA). Quality filtering and read trimming was performed using fastp (version 0.23.2)^50^ at the default settings other than removing reads < 75bp after quality control. Host contaminant sequences were removed by mapping reads to the GRCm39 genome using bowtie2 (version 2.4.2)^51^ in very sensitive mode. Species-level taxonomic profiling was performed using sylph (version 0.5.1)^52^ at the default settings against the pre-built-c200 Genome Taxonomy Database (release 214). Pseudo counts for the species relative abundance was obtained by multiplying the species abundance by the library size after quality and host contaminant filtering and aggregated to higher taxonomic levels using the tax_glom function in the phyloseq package (version 1.41.1)^53^ in R. Shannon diversity was calculated using the estimate_richness function in phyloseq after subsampling the lowest observed read depth (7.5M reads). MaAsLin2 (version 1.6.0) in R was used to identify differential abundant taxa. All analyses were run at the default settings other than min prevalence = 0.2.

### Murine colitis model

3 % DSS was administered in drinking water for 6-7 days, and discontinued when the average weights of the mice reached around 85%. For histologic analyses, sections of colon were fixed in 4% paraformaldehyde, paraffin embedded, sectioned, and stained with hematoxylin and eosin. Disease was scored as follows: (a) weight loss (no change = 0; < 5% =1; 6 – 10% = 2; 11 –20% = 3; > 20% = 4), (b) feces (normal = 0; pasty, semiformed= 2; liquid, sticky, or unable to defecate after 5 min = 4), (c) blood (no blood = 0; visible blood in rectum = 1; visible blood on fur = 2), and (d) general appearance (normal = 0; piloerection = 1; lethargy and piloerection = 2; motionless = 4)^54^.

### Nitrate assay

50ul of sample was mixed with modified Griess reagent (50ul each of vanadium chloride (8mg/ml in 1M HCl), sulfanilamide (2mg/ml in 1M HCl) and naphthyl ethylenediamine dihydrochloride (0.1% w/v in water) for 4hrs (urine) or 16hrs (serum and mucosal scrape extract) for reduction of nitrate to nitrite, and the color reaction with nitrite^34,55^. For preparation of mucosal scrape extract, the scrape of large intestine was homogenized in 100ul PBS, centrifuged at full speed for 10min, the supernatant was collected for analysis. For serum and scrape extract samples, protein was removed prior to the assay by precipitation with 100% EtOH (serum) or 1N HCl (scrape). For quantification of nitrite, absorbance was measured using a micro-plate reader (Biotek Synergy 2) set to 540 nm.

### Flow cytometry

Single cell lamina propria suspensions were obtained by first shaking sections of large intestine in 1 mM EDTA/1 mM DTT at 37°C to remove IECs and intraepithelial cells, then incubated remaining tissues in RPMI with 1 mg/mL collagenase/dispase for 30 min at 37°C with shaking. For intracellular cytokine analyses, cells were stimulated with 50 ng/ml PMA (Sigma-Aldrich), and 500 ng/ml ionomycin (Sigma-Aldrich) in the presence of 10 mg/ml brefeldin A for 3.5 hrs at 37°C. Cells were stained using the following fluorescence-conjugated monoclonal antibodies diluted in FACS buffer (2% FBS, 0.01% Sodium Azide, PBS): FITC anti-CD45.2 (Clone: 104, Invitrogen), perCP-efluor710 anti-CD3 (Clone: 17A2, Invitrogen), APC-eFluor 780 anti-CD4 (Clone: RM4-5, eBioscience), and PE anti-TNFα (Clone: MP6-XT22, eBioscience). Dead cells were excluded with the Fixable Violet Dead Cell Stain Kit (Invitrogen). Intracellular Fixation and Permeabilization Buffer Set (Invitrogen) was used for intracellular staining. Samples were acquired on the BD FACSymphony™ A5 and analyzed with FlowJo Software (Treestar).

### Human intestinal organoids

Intestinal organoids (colonoids) were generated from distal colonic endoscopic biopsy samples. Donors were recruited prospectively as controls for participation in the IBD Biorepository protocol at CCHMC (IRB 2011-2285) and histologic analyses confirmed patients as non-IBD healthy controls. Biopsies were incubated in strip buffer (PBS, 5% FBS, 1 mM EDTA, 1 mM DTT), then incubated in wash buffer (Advanced DMEM, 10% FBS, 2 mM L-glutamate, penicillin-streptomycin) with 2 mg/mL collagenase type I (Invitrogen) at 37°C for 5-15 mins with vigorous mixing as previously described^56^. Crypts were plated in Matrigel with Human Organoid Growth Media (StemCell) supplemented with 10 μM Y-27632, and 10 μM SB 431542. Once organoids were established, they were maintained in human organoid culture media (50% Advanced DMEM/F12 supplemented with 10 mM HEPES, 2 mM L-glutamate, 50% L-WRN conditioned media, 1x N2-supplement, 1x B27-supplement, 50-ng/mL human EGF, 10µM Y-27632, 10 μM SB 431542, and Normocin, 50μg/ml). For experiments, organoids were treated with h-IFNγ (PeproTech), 1,000U/ml and aminoguanidine, 5mM, for 72hrs. Culture supernatant was used for nitrate measurement after de-proteinization with 100% ethanol.

### Statistics

Results are shown as mean ± SEM. Mice of the indicated genotypes were assigned at random to groups. Tests between two groups used a two-tailed Student’s t-test. Survival curves were evaluated by Mantel-Cox test. Results were considered significant at \**p* ≤ 0.05; \*\**p* ≤ 0.01, \*\*\**p* ≤ 0.001. Statistical analyses were performed using Prism version 10.0.

## Acknowledgements

We thank members of the Alenghat lab and Center for Inflammation and Tolerance at CCHMC for useful discussions and critical reading of the manuscript. We thank CCHMC Veterinary Services, Research Flow Cytometry Core, Pathology Research Core, and University of Cincinnati Genomics Core for services and technical assistance. The floxed Nos2 mice were generated by the Morrison and Vazquez-Torres laboratories at the University of Colorado and inquiries about their generation should be directed to them. This research is supported by the National Institutes of Health (DK114123, DK116868, DK137771 to T.A.; K08 DK134884 to S. H. H., R01AI153442 E.R.M.) and a Crohn’s & Colitis Foundation award to T.A. T.A. holds an Investigator in the Pathogenesis of Infectious Disease Award from the Burroughs Wellcome Fund. This project is supported in part by Cincinnati Children’s ARC grant #53671 (E.R.M., L.A.D.), P30 DK078392, and U54 DK126108.

## SUPPLEMENTAL FIGURE LEGENDS

**Supplemental Figure 1. (A)** Phylum level comparison of stool bacterial communities in 8 week old Nos2^FF^ mice and Nos2^ΔIEC^ mice separated by genotype at weaning. **(B)** alpha-diversity as measured by the Shannon Index of samples in (A). Data include n=3 mice per genotype. Results are mean ± SEM.

**Supplemental Figure 2.** Nos2 is specifically elevated and epigenetically altered in intestinal epithelial cells (IECs) isolated from the ileum of IBD patients. Nos2 expression by IECs controls intestinal nitrate levels, sustains intestinal colonization with nitrate-metabolizing bacteria, and increases susceptibility to microbiota-sensitive colitis. Systemic nitrate levels can also serve as a sensitive, non-invasive biomarker for assessing a patient’s intestinal status. Thus, epithelial Nos2 directs host-microbiota dynamics that prime an inflammatory intestinal environment and nitrate metabolism could represent a key pathway to target for monitoring and treating IBD.

## References

1. Wang, R., Li, Z., Liu, S. & Zhang, D. Global, regional and national burden of inflammatory bowel disease in 204 countries and territories from 1990 to 2019: a systematic analysis based on the Global Burden of Disease Study 2019. BMJ Open 13, e065186–e065186 (2023).

2. Dolinger, M., Torres, J. & Vermeire, S. Crohn’s disease. The Lancet 403, 1177–1191 (2024).

3. Alsoud, D., Verstockt, B., Fiocchi, C. & Vermeire, S. Breaking the therapeutic ceiling in drug development in ulcerative colitis. The Lancet Gastroenterology & Hepatology 6, 589–595 (2021).

4. Agrawal, M., et al. Early life exposures and the risk of inflammatory bowel disease: Systematic review and meta-analyses. eClinicalMedicine 36(2021).

5. Rudbaek, J.J., et al. Deciphering the different phases of preclinical inflammatory bowel disease. Nature Reviews Gastroenterology & Hepatology 21, 86–100 (2024).

6. Lloyd-Price, J., et al. Multi-omics of the gut microbial ecosystem in inflammatory bowel diseases. Nature 569, 655–662 (2019).

7. Yilmaz, B., et al. Microbial network disturbances in relapsing refractory Crohn’s disease. Nature Medicine 25, 323–336 (2019).

8. Ryan, F.J., et al. Colonic microbiota is associated with inflammation and host epigenomic alterations in inflammatory bowel disease. Nature Communications 11(2020).

9. Förstermann, U. & Sessa, W.C. Nitric oxide synthases: Regulation and function. in European Heart Journal, Vol. 33 (2012).

10. Aktan, F. iNOS-mediated nitric oxide production and its regulation. Life Sciences 75, 639–653 (2004).

11. Andrabi, S.M., et al. Nitric Oxide: Physiological Functions, Delivery, and Biomedical Applications. Advanced Science 10, 2303259–2303259 (2023).

12. Kang, Y., Liu, R., Wu, J.X. & Chen, L. Structural insights into the mechanism of human soluble guanylate cyclase. Nature 574, 206–210 (2019).

13. Jia, J., et al. Target-selective protein S-nitrosylation by sequence motif recognition. Cell 159, 623–634 (2014).

14. Cross, R.K., Wilson, T. & Wilson, K.T. Nitric Oxide in Inflammatory Bowel Disease. (2003).

15. Kolios, G., Valatas, V. & Ward, S.G. Nitric oxide in inflammatory bowel disease: A universal messenger in an unsolved puzzle. in Immunology, Vol. 113 427–437 (2004).

16. Lundberg, J.O. & Weitzberg, E. Biology of nitrogen oxides in the gastrointestinal tract. Gut 62, 616–629 (2013).

17. Didriksen, B.J., Eshleman, E.M. & Alenghat, T. Epithelial regulation of microbiota-immune cell dynamics. in Mucosal Immunology, Vol. 17 303-313 (Elsevier B.V., 2024).

18. Pool, M.O., et al. Serum nitrate levels in ulcerative colitis and crohn’s disease. Scandinavian Journal of Gastroenterology 30, 784–788 (1995).

19. Goggins, M.G., et al. Increased urinary nitrite, a marker of nitric oxide, in active inflammatory bowel disease. Mediators of Inflammation 10, 69–73 (2001).

20. Murray, I.A., Daniels, I., Coupland, K., Smith, J.A. & Long, R.G. Increased activity and expression of iNOS in human duodenal enterocytes from patients with celiac disease. (2002).

21. Singer, I.I., et al. Expression of Inducible Nitric Oxide Synthase and Nitrotyrosine in Colonic Epithelium in Inflammatory Bowel Disease. in GASTROENTEROLOGY, Vol. 111 871–885 (1996).

22. Gochman, E., et al. The expression of iNOS and nitrotyrosine in colitis and colon cancer in humans. Acta Histochemica 114, 827–835 (2012).

23. Elmentaite, R., et al. Single-Cell Sequencing of Developing Human Gut Reveals Transcriptional Links to Childhood Crohn’s Disease. Developmental Cell 55, 771–783.e775 (2020).

24. Kelly, D., et al. Microbiota-sensitive epigenetic signature predicts inflammation in Crohn’s disease. JCI Insight 3(2018).

25. Nakanishi, Y., Sato, T., Takahashi, K. & Ohteki, T. IFN-γ-dependent epigenetic regulation instructs colitogenic monocyte/macrophage lineage differentiation <em>in vivo</em<. Mucosal Immunology 11, 871–880 (2018).

26. Zhao, X., et al. MEF2C promotes M1 macrophage polarization and Th1 responses. Cellular & Molecular Immunology 19, 540–553 (2022).

27. Hokari, R., et al. Reduced sensitivity of inducible nitric oxide synthase-deficient mice to chronic colitis. Free Radic Biol Med. 31(2):153–63. (2001).

28. Krieglstein, C.F., et al. Regulation of Murine Intestinal Inflammation by Reactive Metabolites of Oxygen and Nitrogen: Divergent Roles of Superoxide and Nitric Oxide. J. Exp. Med.194 1207-1218 (2001).

29. Beck, P.L., et al. Paradoxical roles of different nitric oxide synthase isoforms in colonic injury. Am J Physiol Gastrointest Liver Physiol. 286(1):G137–47. (2004).

30. Forster, S.C., et al. Identification of gut microbial species linked with disease variability in a widely used mouse model of colitis. Nature Microbiology 7, 590–599 (2022).

31. Le Bras, A. Gut microbiota drives disease variability in the DSS colitis mouse model. Lab Animal 51, 131 (2022).

32. Roy, U., et al. Distinct Microbial Communities Trigger Colitis Development upon Intestinal Barrier Damage via Innate or Adaptive Immune Cells. Cell Reports 21, 994–1008 (2017).

33. Rojas-Tapias, D.F., et al. Inflammation-associated nitrate facilitates ectopic colonization of oral bacterium Veillonella parvula in the intestine. Nature Microbiology 7, 1673–1685 (2022).

34. Winter, S.E., et al. Host-Derived Nitrate Boosts Growth of E. coli in the Inflamed Gut. Science 339,708–711(2013).

35. Oberc, A.M., Fiebig-Comyn, A.A., Tsai, C.N., Elhenawy, W. & Coombes, B.K. Antibiotics Potentiate Adherent-Invasive E. coli Infection and Expansion. Inflammatory Bowel Diseases 25, 711–721 (2019).

36. Qi, Q., et al. Hydrogen sulfide produced by the gut microbiota impairs host metabolism via reducing GLP-1 levels in male mice. Nature Metabolism 6, 1601–1615 (2024).

37. Soufli, I., et al. Nitric Oxide, Neutrophil/Lymphocyte, and Platelet/Lymphocyte Ratios as Promising Inflammatory Biomarkers in Complicated Crohn’s Disease: Outcomes of Corticosteroids and Anti-TNF-α Therapies. Inflammation 46, 1091–1105 (2023).

38. Li, J., et al. Identification and multimodal characterization of a specialized epithelial cell type associated with Crohn’s disease. Nature Communications 15, 7204 (2024).

39. Glynne, P.A., Darling, K.E.A., Picot, J. & Evans, T.J. Epithelial inducible nitric-oxide synthase is an apical EBP50-binding protein that directs vectorial nitric oxide output. Journal of Biological Chemistry 277, 33132–33138 (2002).

40. Rumbo, M., Courjault-Gautier, F., Sierro, F., Sirard, J.C. & Felley-Bosco, E. Polarized distribution of inducible nitric oxide synthase regulates activity in intestinal epithelial cells. FEBS Journal 272, 444–453 (2005).

41. Webster, C.M. & Shepherd, M. The nitric oxide paradox: antimicrobial and inhibitor of antibiotic efficacy. Emerging Topics in Life Sciences 8, 37–43 (2023).

42. Hall, J.R., et al. Mode of Nitric Oxide Delivery Affects Antibacterial Action. ACS Biomaterials Science & Engineering 6, 433–441 (2020).

43. Ahmed, N. Advanced glycation endproducts—role in pathology of diabetic complications. Diabetes Research and Clinical Practice 67, 3–21 (2005).

44. Nanki, K., et al. Somatic inflammatory gene mutations in human ulcerative colitis epithelium. Nature 577, 254–259 (2020).

45. Pavlidis, P., et al. Cytokine responsive networks in human colonic epithelial organoids unveil a molecular classification of inflammatory bowel disease. Cell Reports 40, 111439 (2022).

46. Wayman, J.A., et al. Accessible chromatin maps of inflammatory bowel disease intestine nominate cell-type mediators of genetic disease risk. bioRxiv, 2024.2002.2009.579678 (2024).

47. Dobin, A., et al. STAR: ultrafast universal RNA-seq aligner. Bioinformatics 29, 15–21 (2013).

48. Quinlan, A.R. & Hall, I.M. BEDTools: a flexible suite of utilities for comparing genomic features. Bioinformatics 26, 841–842 (2010).

49. Fite, A., et al. Identification and quantitation of mucosal and faecal desulfovibrios using real time polymerase chain reaction. Gut 53, 523 (2004).

50. Chen, S., Zhou, Y., Chen, Y. & Gu, J. fastp: an ultra-fast all-in-one FASTQ preprocessor. Bioinformatics 34, i884–i890 (2018).

51. Langmead, B. & Salzberg, S.L. Fast gapped-read alignment with Bowtie 2. Nature Methods 9, 357–359 (2012).

52. Shaw, J. & Yu, Y.W. Rapid species-level metagenome profiling and containment estimation with sylph. Nature Biotechnology (2024).

53. McMurdie, P.J. & Holmes, S. phyloseq: An R Package for Reproducible Interactive Analysis and Graphics of Microbiome Census Data. PLOS ONE 8, e61217 (2013).

54. Alenghat, T., et al. Histone deacetylase 3 coordinates commensal-bacteria-dependent intestinal homeostasis. Nature 504, 153–153 (2013).

55. Doane, T.A. & Horwáth, W.R. Spectrophotometric Determination of Nitrate with a Single Reagent. Analytical Letters 36, 2713–2722 (2003).

56. VanDussen, K.L., et al. Development of an enhanced human gastrointestinal epithelial culture system to facilitate patient-based assays. Gut 64, 911 (2015).

