## Supplemental Figures for "Epithelial-intrinsic nitric oxide synthase 2 sustains host-microbiota dynamics that promote colitis"

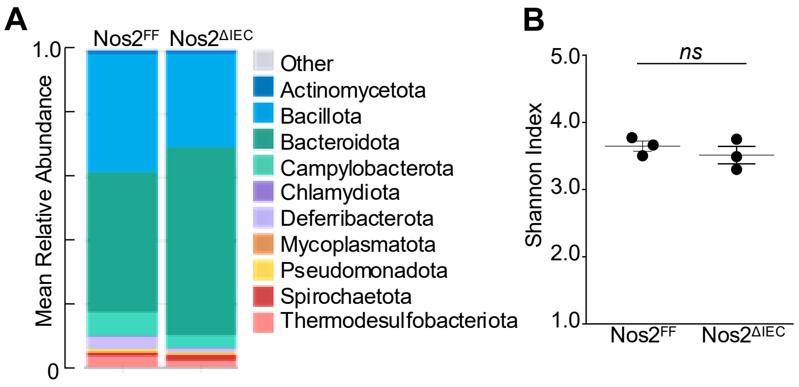


**Supplemental Figure 1. (A)** Phylum level comparison of stool bacterial communities in 8 week old Nos2$^{FF}$ mice and Nos2$^{\Delta IEC}$ mice separated by genotype at weaning and **(B)** alpha-diversity by Shannon Index of samples in (A). Data include n=3 mice per genotype. Results are mean ± SEM.

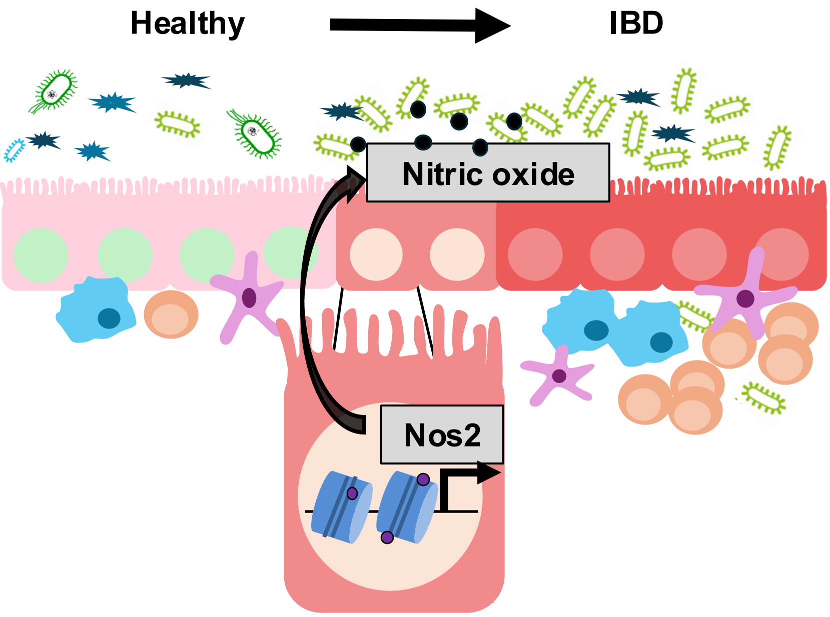


**Supplemental Figure 2.** Nos2 is specifically elevated and epigenetically altered in intestinal epithelial cells (IECs) isolated from the ileum of IBD patients. Nos2 expression by IECs controls intestinal nitrate levels, sustains intestinal colonization with nitrate-metabolizing bacteria, and increases susceptibility to microbiota-sensitive colitis. Systemic nitrate levels can also serve as a sensitive, non-invasive biomarker for assessing a patient's intestinal status. Thus, epithelial Nos2 directs host-microbiota dynamics that prime an inflammatory intestinal environment and nitrate metabolism could represent a key pathway to target for monitoring and treating IBD.
